# Cellular remodeling during photosymbiosis establishment revealed by single-cell dual proteomics

**DOI:** 10.64898/2026.04.21.719821

**Authors:** Chia-Ling Yang, Md Mostafa Kamal, Leandro Ravael, Chi-Yen Wei, Yan-Jun Chen, Sofia Magno, Chin-Wen Chen, Pei-Yi Lin, Chuan-Chih Hsu, Jun-Yi Leu, Chuan Ku

## Abstract

Endosymbiosis of phytoplankton in heterotrophic hosts is ecologically important and has led to key evolutionary innovations. However, the dynamic molecular processes underlying endosymbiosis establishment remain poorly understood. Here, using large-particle sorting and liquid chromatography-tandem mass spectrometry, we unravel heterogeneous changes in proteomes of the cosmopolitan ciliate *Paramecium* and algal endosymbiont *Chlorella* from engulfment to stable endosymbiosis. The initial digestion sees a sharp decline of intracellular *Chlorella* cells, along with host cellular reorganization involving a reduction of the cortex-localized defensive organelles, trichocysts, and proteins for intracellular transport and recycling. The remaining *Chlorella* cells enter a bottleneck stage characterized by energy production and cell cycle commitment before active proliferation. Comparison of *Paramecium* with successful and failed endosymbiosis further identifies a solute carrier transporter that potentially mediates metabolic homeostasis of the endosymbiotic system. Our study reveals inter-organismal coordination during the transition from predator-prey to host-endosymbiont relationships. The approach of time-course single-cell dual proteomics can be useful for investigating diverse interactions between microbial eukaryotes.

## Introduction

Endosymbiosis, a phenomenon where one organism lives inside another, is a major driving force in eukaryote evolution. In addition to the mitochondrial origin, prokaryote-eukaryote and eukaryote-eukaryote endosymbioses have given rise to primary and complex plastids as well as all the photosynthetic lineages of eukaryotes ^1–3^. Algae establish photosymbiotic relationships with diverse heterotrophic hosts, including animals (e.g., cnidarians, molluscs), fungi (lichens), and a wide variety of microbial eukaryotes (e.g., ciliates, rhizarians, amoebozoans) ^4–8^, which play crucial roles in ecosystem functioning by mediating energy and carbon flows ^9,10^. In corals, endodermal cells harbor dinoflagellate algae that provide organic carbon, modulate host physiology, contribute to primary productivity, and facilitated the range expansion of corals ^9,11,12^. Ciliates—one of the most species-rich groups of unicellular predators in both freshwater and marine environments—commonly retain specific algal prey to form mixotrophic holobionts, where hosts and endosymbionts exchange O_2_, CO_2_, organic matter, and other nutrients, and protect each other against competitors, viruses or adverse physical conditions ^5,13–15^. Despite the profound biological impacts of endosymbiosis on unicellular eukaryotes, the molecular changes and cellular remodeling that accompany the transition from predator-prey to host-endosymbiont interactions remain poorly understood.

The ciliate *Paramecium bursaria* (Parameciidae, Oligohymenophorea) and the chlorophyte alga *Chlorella* (Chlorellaceae, Trebouxiophyceae), both widespread in freshwater and other aquatic ecosystems, are commonly used to study endosymbiosis formed by free-living eukaryotes. After encountering *Chlorella*, alga-free (aposymbiotic) *P*. *bursaria* can establish endosymbiosis and harbor hundreds of *Chlorella* cells in its cellular cortex and in close proximity to mitochondria ^16^. This spatial association facilitates the exchange of photosynthetic products and nitrogenous nutrients (e.g., host-derived glutamine), enhancing their survival under thermal and nitrogen-deficiency stress ^17,18^. Microscopic analyses showed that after *P*. *bursaria* phagocytoses *Chlorella* cells, part of the digestive vacuole (DV) transforms into perialgal vacuoles (PV) that surround individual algal cells and prevent host digestion ^19,20,5^. However, the trajectory of endosymbiosis formation, from algal engulfment to the proliferation of intracellular symbionts, has been difficult to capture. The molecular responses of *Chlorella* during this process also remain largely unexplored. To resolve cellular dynamics during endosymbiosis establishment, we need high-resolution, integrative approaches that enable the quantification of molecules in both *Paramecium* and *Chlorella*.

Individual differences are inherent in unicellular organisms, yet such heterogeneity can be obscured by bulk population sampling ^21,22^. Quantifying the abundance of protein species allows us to examine the functions and compositions of cells. Recent advancements in single-cell proteomics (SCP), high-throughput mass spectrometry (MS) analyses of polypeptides from individual cells, have led to the detection of over a thousand protein groups per cell in model mammalian and flowering plant systems ^23,24^. Still, its application to non-model and microbial systems remains limited. The size of *Paramecium* cells (∼50–250 µm) also poses a technial challenge to their single-cell isolation using common fluorescence-activated cell sorters (< 40 µm). In this study, we developed a new SCP platform for large microeukaryotes by integrating a large-particle sorter with data-independent acquisition (DIA)-based LC-MS/MS analyses. This approach enable us to profile protein content dynamics in both the host and endosymbionts during their establishment of endosymbiosis.

## Results

### Endosymbiosis establishment is dynamic and heterogeneous

After mixing (“re-infecting”) aposymbiotic *P*. *bursaria* cells with *Chlorella* cells (originally isolated from stable symbiotic *P*. *bursaria* cells), we removed unengulfed algal cells at 1 hour post-infection (HPI) and sampled the mixed culture throughout a 21-day time-course experiment at 3 HPI, 1 day post-infection (DPI), 2 DPI, 4 DPI, 7 DPI, 14 DPI, and 21 DPI. The chlorophyll a fluorescence intensity per *P. bursaria* cell was quantified using a COPAS Vision large-particle flow cytometer with 488-nm excitation. Intracellular *Chlorella* cells were also manually counted under fluorescence microscopes. At 3 HPI, the vast majority of *P. bursaria* cells had higher chlorophyll fluorescence than the aposymbiotic control (Fig. 1A), with > 90% of *P. bursaria* cells containing > 20 *Chlorella* cells and nearly 60% having > 100 *Chlorella* cells (Fig. 1B-F). We also observed a few yellow-green or brown algal cells, suggesting they were enclosed by DVs and undergoing digestion at 3 HPI (Fig. 1E). The *Chlorella* cell counts decreased from 3 HPI to 2 DPI, when 95% of the *P. bursaria* contained ≤ 20 *Chlorella* cells. Since 2 DPI, the proportion of *P. bursaria* cells housing > 100 *Chlorella* cells continually increased, with 3% at 4 DPI, 65.8% at 7 DPI, and 100% at 21 DPI (Fig. 1B). It indicates that intracellular *Chlorella* began to actively divide around 2–4 DPI. Newly proliferated *Chlorella* cells (e.g., Fig. 1I, J) had a more peripheral distribution distinct from the engulfed cells at 3 HPI (Fig. 1E, F). The *P. bursaria* population exhibited a bimodal distribution at 7 and 14 DPI, with most cells having either high (> 100) or low (≤ 20) *Chlorella* counts (Fig. 1B and G-J). Similar patterns can be observed in the distribution of chlorophyll fluorescence per *P. bursaria* cell, where cells with low or no fluorescence can still be seen at 7 and 14 DPI (Fig. 1A). Under the high light condition (21.8 μmol·m^−2^·s^−1^), low-chlorophyll *P. bursaria* cells, which were similar to alga-free cells and thus less tolerant to light ^13^, gradually died out and were not observed anymore at 21 DPI (Fig. 1A, B). These observations indicate that the establishment of *Paramecium*-*Chlorella* endosymbiosis—from phagocytosis and digestion to proliferation of endosymbionts and stable endosymbiosis—is a highly dynamic and heterogeneous process at the level of individual cells.

**Figure 1.**
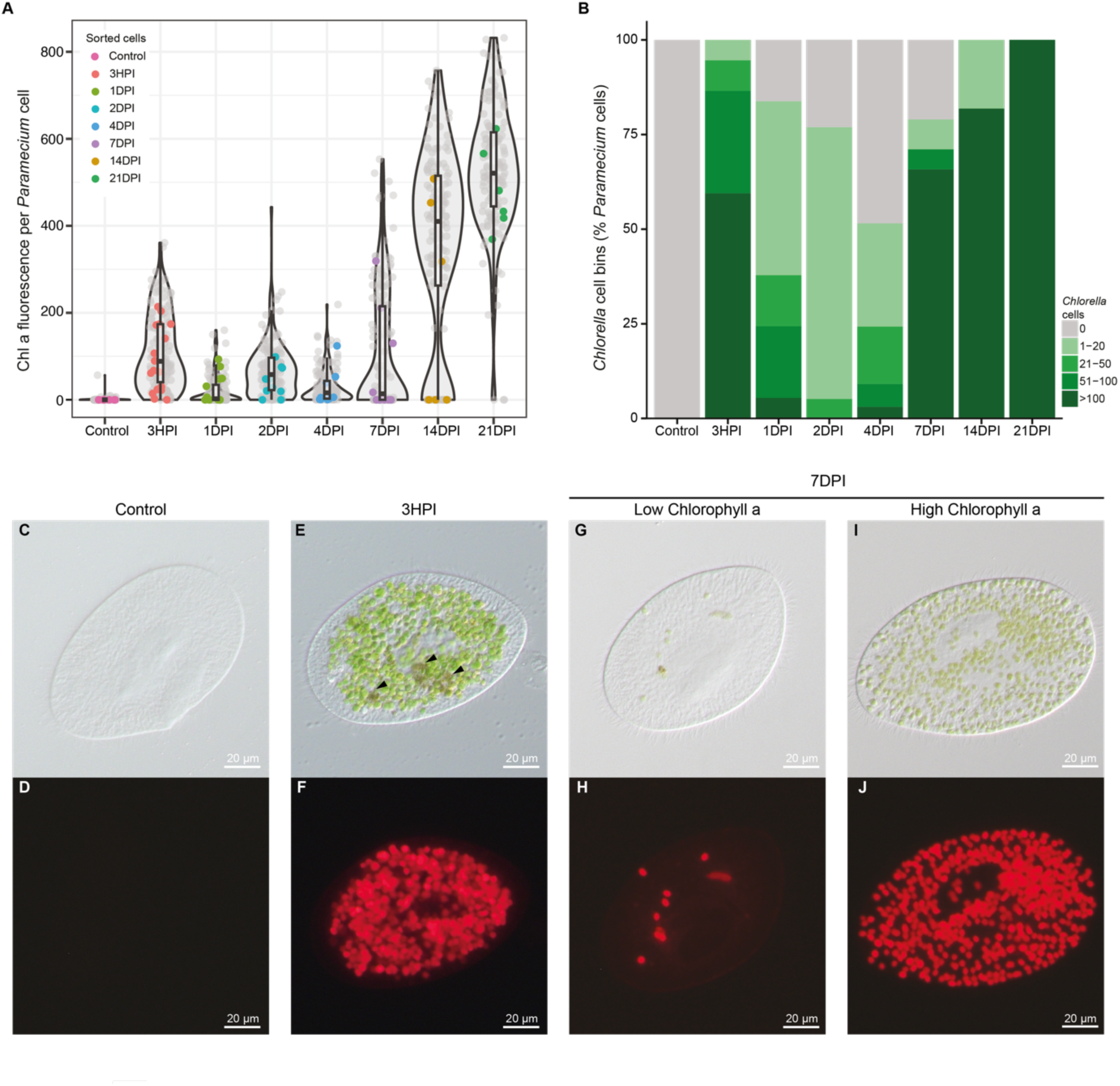
Intracellular algal loads in *Paramecium* cells throughout the endosymbiosis establishment. (A) Distribution of fluorescence intensities (pulse area at 680/42 nm; a.u.) corresponding to chlorophyll a abundance of individual *P. bursaria* cells measured through flow cytometry. Each dot represents a single cell, with colored dots indicating the randomly isolated cells for SCP analyses (n=6, 18, 11, 10, 9, 10, 6, 6). Aposymbiotic cells are used as the control. (B) Distribution of *Chlorella* cell counts within individual *P. bursaria* cells from each sample (n=30, 37, 37, 36, 32, 38, 33, 40). (C-J) Representative brightfield and fluorescence micrographs of *P. bursaria* cells that are aposymbiotic (control), at 3 HPI, or at 7 DPI. The abundance and distribution of *Chlorella* cells vary at the same timepoint and between timepoints. (E) Digested (arrowheads) *Chlorella* cells can be observed at 3 HPI.

### Three distinct stages of endosymbiosis establishment

To further understand the cellular changes during endosymbiosis establishment, we randomly sorted individual *P. bursaria* cells at each time point (Fig. 1A) and profiled the single-cell proteomes of 80 cells (Supplementary Table S1). We analyzed DIA MS spectra with Spectronaut (Supplementary Table S2) using a combined database of predicted sequences from *P. bursaria* and *Chlorella* (Supplementary Table S3) to identify protein groups (i.e., clusters of protein isoforms or homologs that share identical peptide sequences). After filtering out lowly represented protein groups, we identified 1599 *P. bursaria* protein groups (mean±SD per cell: 1284.1±194.3) and 231 *Chlorella* protein groups (per cell: 91.4±71.4) (Fig. 2A, B). Four cells with < 100 protein groups were excluded from further analyses.

**Figure 2.**
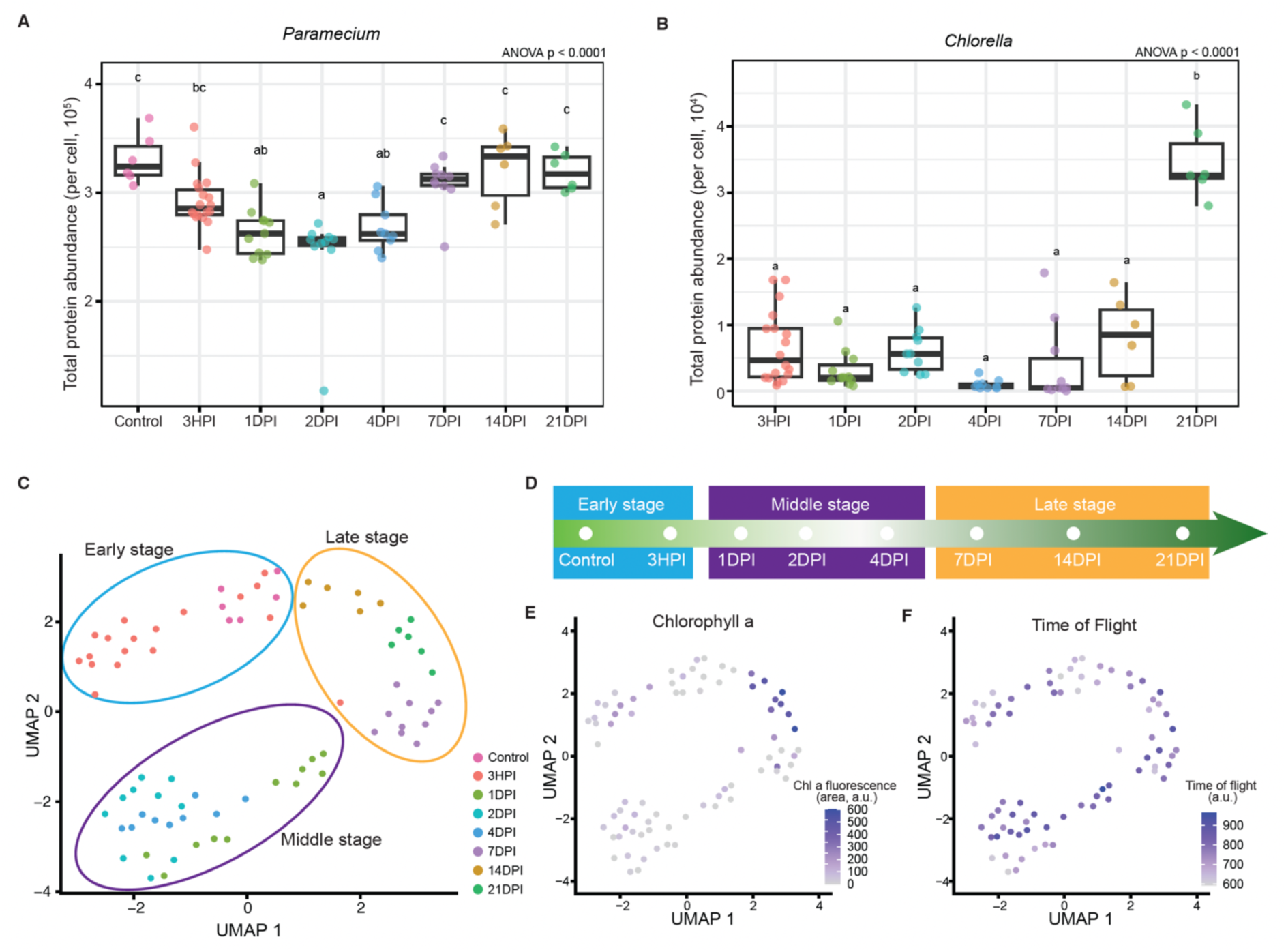
Single-cell proteomic analysis of *P. bursaria* during the formation of endosymbiosis with *Chlorella*. (A, B) Total abundance (sum of peptide fragment signal areas) of *Paramecium*-encoded (A) and *Chlorella*-encoded (B) proteins detected in individual *P. bursaria* cells. (C) UMAP projection of SCP profiles of *Paramecium*-encoded proteins and grouping of cells into the early, middle, and late stages of endosymbiosis establishment (D). (E, F) Mapping of fluorescence (680/42 nm) for chlorophyll a abundance (E; Fig. 1A) and time of flight (TOF) for cell size (F) onto the UMAP projection. (A, B) One-way ANOVA with correction by Tukey’s HSD test.

The total abundance of *P. bursaria* (Fig. 2A) and *Chlorella* (Fig. 2B) proteins detected (sum of peptide fragment ion intensities) in individual *P. bursaria* cells shows distinct changes through time, with a strong correlation (R^2^ = 0.81) between total *Chlorella* protein abundance and chlorophyll fluorescence (Extended Data Fig. 1). The *P. bursaria* protein abundance was significantly reduced from the aposymbiotic state to 2 DPI by ∼25%, then it increased back to the original levels from 2 DPI to 21 DPI. The *Chlorella*-encoded proteins decreased from 3 HPI to 4 DPI, then substantially increased in some cells at 7 and 14 DPI, and reached an average level at 21 DPI that was ∼6 times higher than at 3 HPI.

For further analyses, the SCP data of each cell were normalized and log-transformed using the LogNormalize method in Seurat ^25^ to account for the sparsity and technical noise in single-cell datasets. We performed dimensional reduction with Uniform Manifold Approximation and Projection (UMAP)^26^ using the 221 most variable *P. bursaria* protein groups (Seurat selection method: mean.var.plot). The UMAP distribution of the cells (Fig. 2C) largely aligns with the early (aposymbiotic state and 3 HPI), middle (1, 2, and 4 DPI), and late (7, 14, and 21 DPI) sampling time points of the cells (Fig. 2D). Mapping chlorophyll fluorescence levels shows that some late-stage cells (all 21-DPI and part of 14- and 7-DPI cells) contained the highest chlorophyll content (Fig. 2E; see also Fig. 1A). Together with the overall increase in *Chlorella* cell counts (Fig. 1B) and total protein abundance (Fig. 2B) from 4 to 21 DPI, these results indicate that those late-stage cells with high chlorophyll content had successfully established an endosymbiotic relationship. The time of flight (indicator of particle size) of sorted cells shows no specific distribution pattern across different stages (Fig. 2F; Extended Data Fig. 1), indicating the reduced *P. bursaria* proteome size in the middle stage was not due to reduction in cell size.

### Proteomic remodeling of Paramecium during the transition from feeding to endosymbiosis

To investigate host proteome reduction in the middle stage, we identified 62 *P. bursaria* proteins that were at least two-fold lower (Wilcoxon rank-sum test: *p* < 0.05) at 2 DPI compared with the early- and late-stage cells combined (Fig. 3A). Among these 2DPI-lower proteins, we note the presence of two trichocyst matrix proteins (TMPs), which comprise trichocysts, the projectile defensive organelles in the cellular cortex of *Paramecium* and other ciliates ^27^. These two TMPs belong to two functionally distinct subfamilies, T4 and T1, which respectively comprise the core and cortex layers of the *Paramecium* trichocyst ^28^. By comparing the aposymbiotic control and 2-DPI cells, we observe an overall ∼2.5-fold change in the average total abundance of the 29 detected TMP protein groups (Fig. 3B; Supplementary Table S4). This suggests that before endosymbiotic *Chlorella* cells populate the host cell cortex in the late stage, the host has already begun to scale down its defensive organelles and potentially reserve cortical space for the endosymbionts to come.

**Figure 3.**
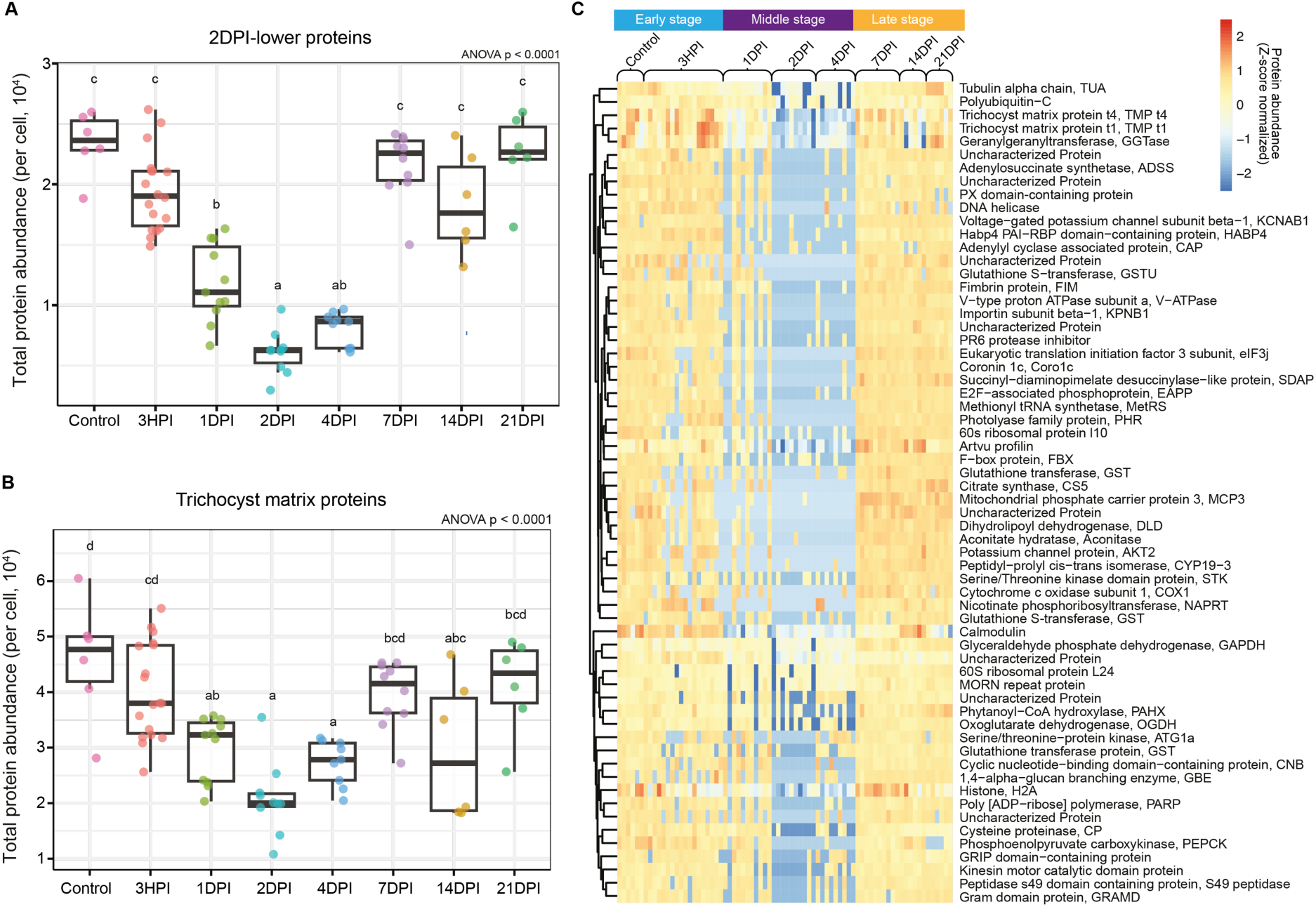
*P. bursaria* proteome remodeling before the proliferation of endosymbiotic *Chlorella*. (A) Total abundance of 2DPI-lower proteins, *Paramecium*-encoded proteins that have significantly lower levels (Wilcoxon rank-sum test: *p* < 0.05) at 2 DPI than the early (aposymbiotic control and 3 HPI) and late (7 DPI, 14 DPI, and 21 DPI) timepoints combined. (B) Total abundance of all detected trichocyst matrix proteins (TMPs) in individual *P. bursaria* cells across samples. (C) Normalized abundance profiles and annotations of the 62 2DPI-lower protein groups. (A, B) One-way ANOVA with correction by Tukey’s HSD test.

The other 2DPI-lower proteins (Fig. 3C; Supplementary Table S5; Extended Data Fig. 2) include a plethora of components in the cytoskeleton and intracellular transport: coronin, fimbrin, profilin, and adenylyl cyclase-associated protein (regulators of actin dynamics); alpha-tubulin, kinesin, and importin (cytoplasm-to-nucleus transport); geranylgeranyltransferase II (modifier of Rab); and MORN repeat and PX domain-containing proteins (regulators of membrane targeting). They also include ATG1, which initiates autophagy to digest cellular components, and vacuolar-type ATPase subunit a, which is part of the V0 domain that mediates proton transport across membranes and acidifies organelles such as lysosomes. Five proteins are involved in redox regulation: four glutathione transferases and nicotinate phosphoribosyltransferase (biosynthesis of NAD). Furthermore, many 2DPI-lower proteins participate in carbohydrate and energy metabolism: glyceraldehyde 3-phosphate dehydrogenase (glycolysis); citrate synthase, aconitase, oxoglutarate dehydrogenase, and dihydrolipoyl dehydrogenase (TCA cycle); mitochondrion-encoded cytochrome c oxidase subunit 1 and mitochondrial phosphate carrier protein (oxidative phosphorylation); and phosphoenolpyruvate carboxykinase (gluconeogenesis). To summarize, the transition from the early stage (engulfment and digestion of most *Chlorella* cells) to the middle stage (remaining *Chlorella* cells starting to proliferate) is marked by a substantial reduction in trichocysts, intracellular transport, membrane trafficking, degradation of cellular components, and energy metabolism.

### Late-stage divergence of symbiotic status and cellular proteomes

*P. bursaria* cells with high numbers of endosymbionts in the late stage all arose from those with few *Chlorella* cells in the middle stage (Fig. 1B); however, not every *P. bursaria* cell reached the highly symbiotic state, as evidenced by the bimodal distribution of algal loads at 7 and 14 DPI (Fig. 1A, B). Our SCP data encompass both cells with high (intensity >100; high-chl) and low chlorophyll levels (intensity ≦ 100; low-chl) (Fig. 1A). *P. bursaria* cells with more intracellular algae appear more opaque in brightfield images collected during sorting and show chlorophyll fluorescence signals along the entire profile of each cell, in sharp contrast to low-chl cells (Fig. 4A). The sorted high- and low-chl cells at 14 DPI had even more pronounced differences in chlorophyll levels (Fig. 1A and Fig. 4A). The 14-DPI low-chl cells are more similar to the aposymbiotic cells and 3-HPI cells (Fig. 2C, E), whereas high-chl cells at 7 and 14 DPI share similar proteomes (Fig. 4B). Together with the low *Chlorella* protein abundances (see below), these findings suggest that late-stage low-chl *P. bursaria* cells and, if any, their remaining intracellular *Chlorella* (e.g., Fig. 1G, H) have failed to establish endosymbiosis. Almost the entire 21-DPI population consists of high-chl cells, with even higher algal loads than at 14 DPI (Fig. 1A, B). Since it took more than 7 days for some of the 4-DPI cells to become high-chl cells at 14 DPI, the absence of the low-chl cells at 21 DPI suggests that those cells that failed to establish endosymbiosis with proliferative *Chlorella* cells eventually died out in the population.

**Figure 4.**
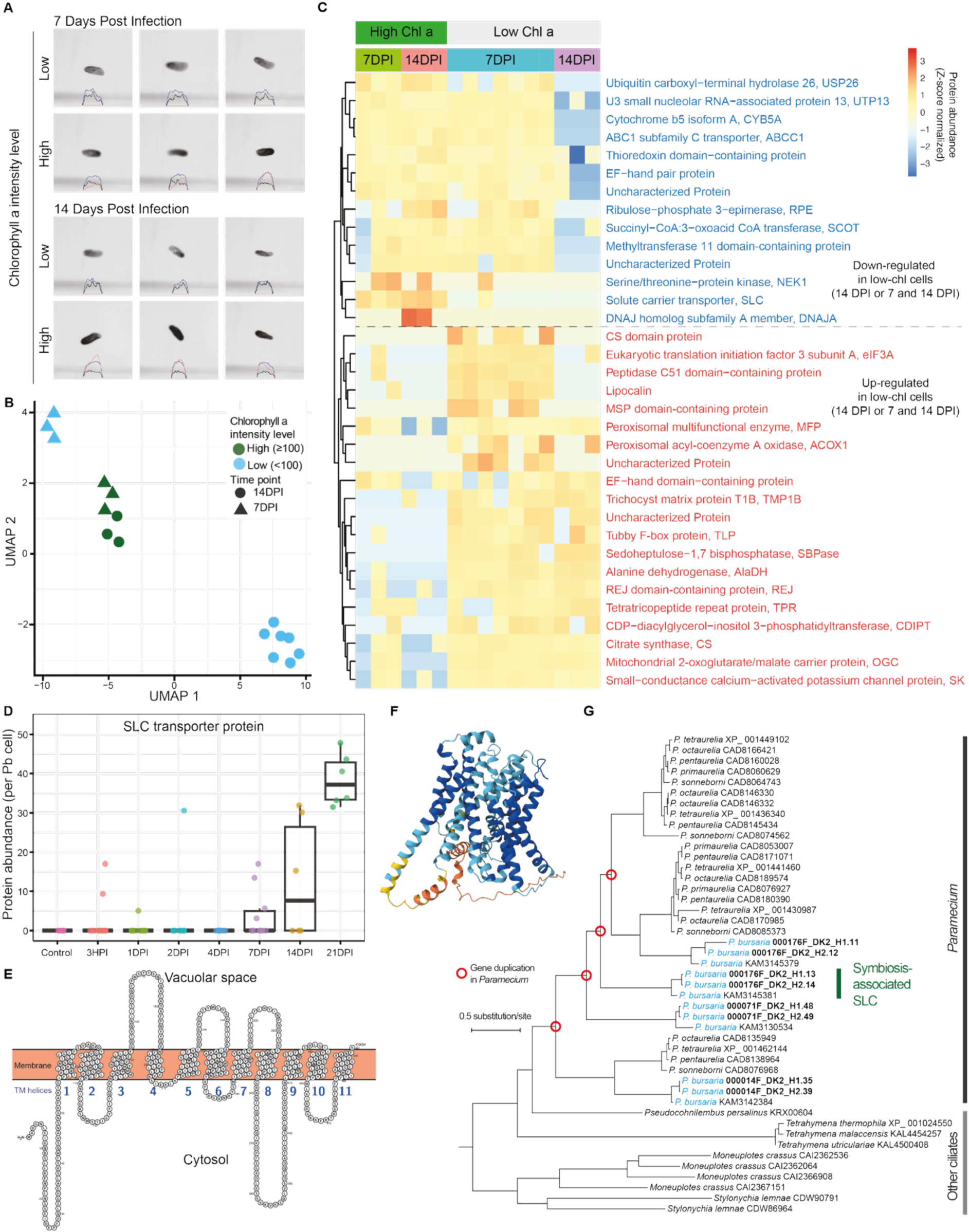
Within-population divergence between *P. bursaria* cells with and without *Chlorella* proliferation. (A) Brightfield images of representative cells with low (< 100) and high (≥ 100) chlorophyll a fluorescence intensities (pulse area at 680/42 nm; a.u.) sorted at 7 and 14 DPI. The red, blue, and black optical profiles of each cell show chlorophyll a fluorescence (680/42 nm), extinction, and forward scatter. (B) Projection of all 7- and 14-DPI cells onto the UMAP 2D space (extracted from Fig. 2C). (C) Normalized abundance profiles and annotations of proteins that are differentially abundant (fold change > 2 or < -2; Wilcoxon rank-sum test: *p* < 0.1) between high and low chlorophyll cells at 14 DPI or 7 and 14 DPI (Supplementary Table S6). (D) Abundance of the SLC transporter proteins in individual *P. bursaria* cells across samples. (E) Predicted topology of the SLC transporter (000176F_DK2_H1.13), showing the cytosolic N-terminus and 11 transmembrane (TM) helices. (F) Predicted structure (pTM = 0.81). Blue (pLDDT: > 90) and light blue (pLDDT: 70-90) indicate high-confidence residues. (G) Maximum likelihood phylogeny of SLC proteins in *Paramecium* and other ciliates. *P*. *bursaria* (light blue) has four proteins as a result of gene duplications within *Paramecium*. The symbiosis-associated SLC protein is unique to *P*. *bursaria*, with its ortholog lost in other, nonsymbiotic *Paramecium* species.

To investigate the physiology associated with different outcomes of endosymbiosis establishment, we extracted proteins with a log2 fold change > 2 or < -2 (Wilcoxon rank-sum test: *p* < 0.1) between high- and low-chl cells at either 14 DPI or at both 7 and 14 DPI (Fig. 4C; Supplementary Table S6; Extended Data Fig. 3). Proteins with higher abundance in high-chl cells at both 7 and 14 DPI include a solute carrier (SLC) transporter protein. The low-chl 14-DPI cells have other proteins with lower abundance: ubiquitin carboxyl-terminal hydrolase 26 (USP26; deubiquitination), U3 small nucleolar RNA-associated protein 13 (UTP13; rRNA maturation), cytochrome b5 (lipid biosynthesis), ABC1 subfamily C transporter (lipid transport), ribulose-5-phosphate 3-epimerase (pentose phosphate pathway), and methyltransferase 11 domain-containing protein (methylation). Overall, the low-chl cells at 14 DPI have lower levels of proteins involved in maintaining biomolecule synthesis and homeostasis.

Proteins with higher abundance in low-chl cells at 7 and 14 DPI include several carbohydrate metabolic enzymes: fructose-1,6-bisphosphatase (or sedoheptulose-1,7-bisphosphatase; gluconeogenesis), alanine dehydrogenase (conversion of alanine to pyruvate), citrate synthase (TCA cycle), and mitochondrial 2-oxoglutarate/malate carrier protein (malate-aspartate shuttle and gluconeogenesis). Two other proteins that are more abundant in low-chl cells are peroxisomal multifunctional enzyme and peroxisomal acyl-CoA oxidase (β-oxidation pathway for breaking down fatty acids and producing energy). These findings suggest that the low-chl cells in the late stage are under starvation and may experience redox stress, consistent with their lack of algae to provide carbohydrates and protection from excessive light. These metabolic enzymes are also among the differentially abundant proteins between 21-DPI cells (all high-chl) and aposymbiotic control cells (lower light condition) (Extended Data Fig. 4). Additional 21-DPI up-regulated proteins include glutamine synthetase (a potential nitrogen source), beta-glucosidase, Complex I subunits, and various proteins for membrane trafficking. Additional aposymbiotic up-regulated proteins contain glutamate dehydrogenase, phosphoenolpyruvate carboxykinase, homogentisate 1,2-dioxygenase (energy production), and SmD3 (spliceosome component).

Compared with 7 and 14 DPI, the SLC transporter further increased at 21 DPI (Fig. 4D), when endosymbiotic *Chlorella* cells are more abundant. The SLC protein has two allelic variants in the *P. bursaria* DK 2 genome (000176F_DK2_H1.13 and 000176F_DK2_H2.14: 96% amino acid identity). A BLAST ^29^ search against the NCBI non-redundant (nr) database ^30^ detected its homologs in other *Paramecium* species and other ciliates, with a sequence identity of ∼45% to the most similar hits (*P*. *octaurelia* and *P*. *pentaurelia*; Supplementary Table S7). Distant homologs are also found in animals (e.g., SLC36A1 [proton-coupled amino acid transporter 1] in mammals) and fungi (e.g., Avt7p in budding yeast). Structural ^31^ and topological ^32^ predictions indicate that the *P. bursaria* SLC has 11 transmembrane helices and a cytosolic N-terminus (Fig. 4E, F). SLCs transport a wide range of substrates, including amino acids, oligopeptides, sugars, nucleotides, lipids, and ions ^33,34^. In addition to plasma membrane, SLCs can be localized to lysosomes and transport amino acids and other breakdown products into the cytoplasm ^35,36^. Subcellular localization prediction by DeepLoc 2.0 ^37^ indicates that the *P. bursaria* SLC most likely localizes to lysosomes or vacuoles (probabilities for the two variants: 0.6339 and 0.6381). We confirm the correctness of DeepLoc 2.0 predictions for well-studied homologs (Supplementary Table S7; Extended Data Fig. 5), including human SLC36A1 ^35^ and yeast Avt7p ^38^, which transport amino acids from either the extracellular and lysosomal space or vacuolar space into cytosol (N-terminal side). Together with the correlation between the abundance of the host-encoded SLC and endosymbiotic *Chlorella* cells, our results suggest that this symbiosis-associated transporter may localize to the perialgal vacuolar membrane of each endosymbiont and maintain metabolic homeostasis by transporting amino acids, sugars, or other nutrients from the endosymbiont into the host cytosol. It is notable that this SLC family underwent four duplications in *Paramecium* before species diversification, and due to loss in the non-symbiotic lineage, the symbiosis-associated SLC is uniquely found in *P*. *bursaria* (Fig. 4G).

### A symbiont bottleneck during endosymbiosis establishment

The abundance of individual *Chlorella* proteins shows distinct patterns across time points (Fig. 5; Supplementary Table S8). Hierarchical clustering of proteomic profiles assigned most Chlorella proteins to 6 classes. Class 1 proteins, with higher relative levels at 2 DPI than 21 DPI, are encoded by genes mainly located in the chloroplast genome. Class 6 proteins similarly have higher levels in the middle stage than the others. Together, these two classes encompass chloroplast- and nucleus-encoded proteins mainly involved in the light-dependent reactions of photosynthesis: Photosystem I (PsaA and PsaB), Photosystem II (PsbA, PsbB, PsbC, PsbD, PsbE, and PsbO), cytochrome *b*_6_*f* complex (PetA and PetB), and ATP synthase α, β, and γ subunits (AtpA, AtpB, and AtpC). However, Calvin cycle proteins are relatively less abundant. It suggests that middle-stage *Chlorella* cells may carry out partial photosynthetic functions to produce ATP and NADPH for use in physiological processes other than carbon fixation.

**Figure 5.**
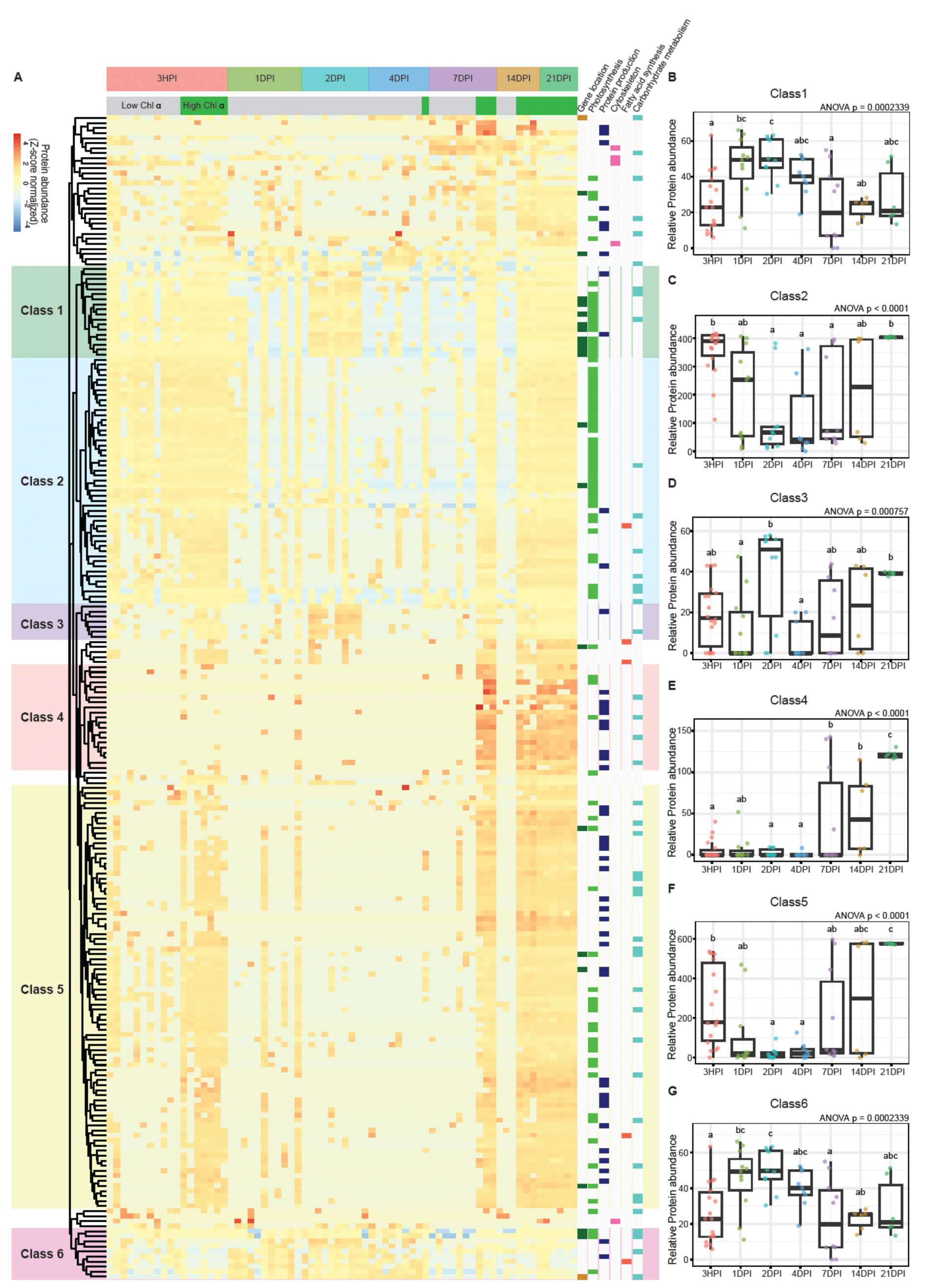
*Chlorella* proteomic changes during the formation of endosymbiosis with *P. bursaria*. (A) Normalized abundance profiles of *Chlorella* proteins ordered through hierarchical clustering. Most detected proteins are assigned into 6 classes based on hierarchical clustering of their abundance patterns across the samples (Supplementary Table S8). Gene location: chloroplast genome (green), mitochondrial genome (brown), nuclear genome (white). (B-G) Total abundance per cell of protein Classes 1–6 in individual *P. bursaria* cells across samples (one-way ANOVA with correction by Tukey’s HSD test).

Other nucleus-encoded proteins with higher abundance at 2 DPI or in the middle stage (Classes 1, 3, and 6) include three histone proteins (H2B, H3, and H4), cell division cycle protein 48 (CDC48), and a cyclin-dependent kinase (CDK), suggesting a potential role of cell cycle progression in shaping the 2-DPI proteomes. Histone abundance varies throughout the cell cycle, with histone accumulation being coupled to cell cycle commitment in eukaryotes^39^. Unlike most eukaryotic organisms, *Chlorella* and some other green algal cells can divide by multiple fission and might have a multinuclear or polyploid stage before division ^40^, which can increase the proportion of histone proteins in the proteome. We also note that two proteins involved in energy metabolism are relatively abundant in middle-stage cells (Class 6). Adenylate kinase maintains cellular energy homeostasis and redistributes energy through the interconversion of ATP, ADP, and AMP. The Δ3,5-Δ2,4-dienoyl-CoA isomerase facilitates β-oxidation of polyunsaturated fatty acids with double bonds at odd positions, such as the major fatty acid in *Chlorella*, α-linolenic acid (C18:3 ω3) ^41^, to generate additional acetyl-CoA for energy production or biosynthesis.

*Chlorella* proteins that are more abundant in the early or late stage (Class 2) are largely additional photosynthesis-related proteins, including reaction center subunits and light-harvesting complex chlorophyll-binding proteins in Photosystems I and II, the ferredoxin-thioredoxin system, RuBisCO subunits, glyceraldehyde-3-phosphate dehydrogenase, phosphoglycerate kinase, Calvin cycle protein CP12, and two low-CO_2_ inducible proteins (potential carbon-concentrating mechanisms). In addition, the presence of granule-bound starch synthase and an acyl carrier protein for fatty acid synthesis indicates that the *Chlorella* cells (freshly engulfed or in the late stage) have the protein complement to actively carry out photosynthesis and convert its products into storage molecules.

Freshly engulfed *Chlorella* cells and the proliferating cells that have just established endosymbiosis are distinguished by Class 4 (almost absent at 3 HPI) and Class 5 (less abundant at 3 HPI) proteins, which predominantly function in translation (ribosomal proteins), photosynthesis and chlorophyll synthesis (RuBisCO activase, geranylgeranyl reductase, protochlorophyllide reductase), and amino acid synthesis (methionine synthase, argininosuccinate synthase, 2-oxoglutarate/malate translocator). Class 4 proteins also include nucleoside diphosphate kinase (nucleotide synthesis), DEAD-box ATP-dependent RNA helicase (RNA metabolism), S-adenosylmethionine synthase (methyl group metabolism), mitochondrial ATP/ADP carrier ANT1 (ATP transport), phosphoglucomutase (starch synthesis), and THYLAKOID FORMATION 1 (thylakoid membrane biogenesis). Altogether, it suggests that after the middle-stage bottleneck, late-stage *Chlorella* cells accumulate more proteins than in the early stage, enabling them to populate *P. bursaria* cells through active growth and proliferation. Conversely, although low-chl *P. bursaria* cells in the late stage may still contain intracellular *Chlorella* (e.g., Fig. 1H), these algal cells have lower protein abundances than those in the early or middle stage (Fig. 5A), suggesting that the remaining *Chlorella* cells in low-chl *P. bursaria* are likely metabolically inactive and already incapable of forming stable symbiosis.

## Discussion

Based on the proteomic, flow cytometry, and microscopy data, we propose a temporal model for endosymbiosis formation between *P. bursaria* and *Chlorella* (Fig. 6). After their encounter, *Chlorella* cells are phagocytosed and mostly digested within 1 day, while *Paramecium* cells enter proteomic remodeling that results in an overall reduction in their proteomes. A decrease in proteins for trichocyst formation, intracellular transport, autophagy, and lysosomal activity, likely reduces energy demand, frees up cortical space, and creates an intracellular environment more conducive to the conversion of food algae into endosymbionts. The surviving, bottlenecked *Chlorella* population is now characterized by increased proteins for energy production (light-dependent reactions and β-oxidation) and a potential commitment to cell cycle progression. After 4 days, *P. bursaria* starts to diverge into a high-chl subgroup that has established endosymbiosis with proliferating *Chlorella* cells and a low-chl one that has no or few remaining *Chlorella* cells which lost their protein machinery for photosynthesis. Without *Chlorella* for food production and photoprotection, the low-chl *P. bursaria* shows signs of starvation and eventually dies out in the population. *P. bursaria* cells with more *Chlorella* endosymbionts express higher levels of an SLC transporter that likely localizes to PVs and transports amino acids or other nutrients into host cytosol. With successful endosymbiosis establishment, intracellular *Chlorella* cells rapidly grow and proliferate, accumulating more proteins for biosynthesis (e.g., amino acids, chlorophyll, starch, and nucleotides). Overall, this study demonstrates how the dynamic dual proteomic remodeling of structural components, metabolic enzymes, and transporter proteins can transform *Paramecium* and *Chlorella* from predator-prey to host-endosymbiont relationships.

**Figure 6.**
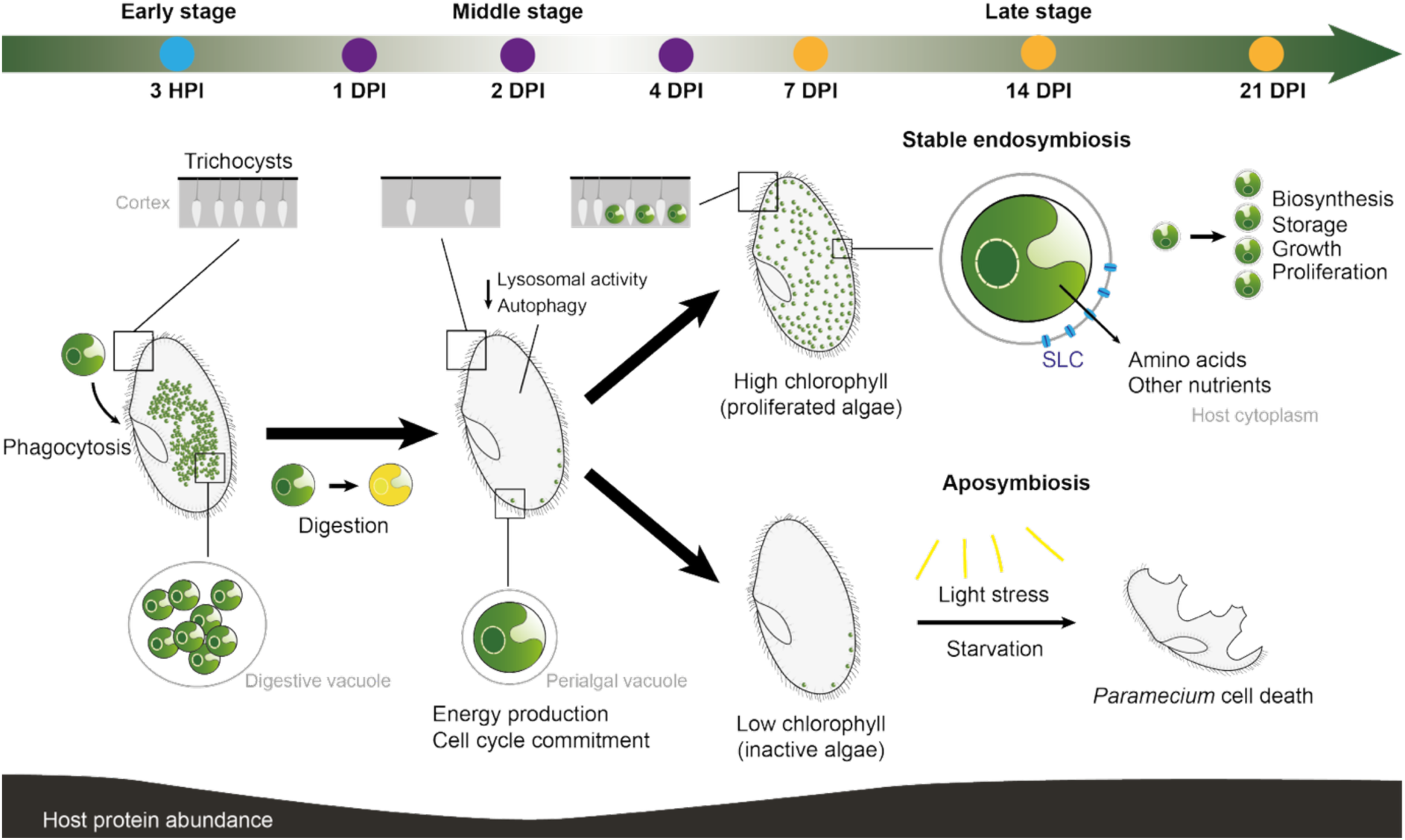
Dynamic process of endosymbiosis establishment between *P. bursaria* and *Chlorella*. Both symbiotic partners undergo proteomic, functional, and cell biological changes from the engulfment of algae to the establishment of algal endosymbionts inside *P. bursaria* cells. Under the high-light growth condition with limited food, aposymbiotic *P. bursaria* cells are eventually replaced by symbiotic ones. Organic compounds synthesized by the endosymbiotic algae can be transported from within perialgal vacuoles to the host cytoplasm, potentially through SLC transporter proteins.

Energy metabolism and nutrient exchange are key to the formation and maintenance of symbiosis ^5,8^. Here, we show that metabolic states of both hosts and endosymbionts change through the different stages of endosymbiosis establishment. During the middle stage, host energy production declines (Fig. 3), whereas *Chlorella* cells harbor more proteins for energy production (Fig. 5). The bioenergetic states of late-stage *P. bursaria* cells depend on the state of endosymbiosis (Fig. 4). With successful endosymbiosis, *P. bursaria* cells contain an increasing number of algal endosymbionts that have a full spectrum of proteins for photosynthesis and translation, as well as biosynthetic enzymes for amino acids and starch (Fig. 5).

One common theme for both symbiotic partners is that proteomic changes may prepare the cells for the next stage of endosymbiosis establishment. It is a well-known observation that *Chlorella* endosymbionts occupy the same cortex space as the native host organelles, trichocysts ^5^. Consistent with the observation that lysosomal fusion may cause digestion of trichocysts after *P*. *bursaria* engulfs *Chlorella* ^42^, SCP profiles reveal a decrease of trichocyst proteins from the early to middle stage. However, there is also an increase of trichocyst proteins from the middle to late stage (Fig. 3B), suggesting *P*. *bursaria* temporarily reduce trichocysts for cellular remodeling during endosymbiosis establishment. Based on the results, we propose a destroy-and-rebuild model for the cortex space remodeling (Fig. 6). In the “destroy” phase, the decline of trichocysts, which precedes *Chlorella* proliferation, is not due to direct competition from or replacement by *Chlorella* cells but mainly caused by lysosomal fusion. This low-trichocyst cortex environment may be more suitable for *Chlorella* invasion and agrees with the finding that trichocysts are unnecessary for endosymbiont localization ^16^. In the “rebuild” phase, trichocysts are regenerated alongside proliferating *Chlorella* cells, resulting in their compact arrangement in symbiotic *P*. *bursaria* cells.

The rapid increase of *Chlorella* cells from < 20 at 2 DPI to > 100 at 7 DPI in most *P. bursaria* cells (Fig. 1B) may be attributable to bottlenecked *Chlorella* cells having more proteins for energy production, histones, and regulators of cell cycle progression (Fig. 5A; Supplementary Table S8). A larger proportion of histones in the proteome is consistent with the occurrence of multiple fissions in endosymbiotic *Chlorella* ^18^. It also suggests that the cells are already committed to cell division or already divide into daughter cells within the shared mother cell wall, which can account for the rapid cell proliferation that follows. In contrast to the middle-stage *Chlorella* cells that prioritize reproduction for colonizing the host cortex, established endosymbionts in the late stage invest more resources on growth and storage.

It has been suggested that *Chlorella* endosymbionts provide O_2_, maltose, and other organic compounds to *Paramecium* hosts and, in return, receive CO_2_ and nitrogen in glutamine or other forms ^5,17,43^. However, the specific transporters or mechanisms for moving these nutrients between endosymbionts and hosts remain unknown. Our SCP data and protein localization prediction reveal a host-encoded SLC as a potential perialgal vacuole-localized transporter. SLCs commonly facilitate nutrient influx into cytosol and can be uniporters, cotransporters (e.g., sodium-glucose cotransporters), or antiporters (e.g., SLC7A11 that exchanges cystine with glutamate) ^33^. The same SLC can be found at different subcellular locations and be promiscuous in substrate selection ^44^. While the *P. bursaria* symbiosis-associated SLC likely contributes to host nutrient intake, it is also possible that it functions as an antiporter, transporting nutrients (e.g., glutamine) into the perialgal vacuole as well. The importance of SLCs in mediating host-endosymbiont metabolite exchange is well known for mitochondria ^45^ and animal-microbe symbiosis ^46^. Our study indicates that they may also play a key role in establishing algal endosymbiosis in *P. bursaria* and other eukaryotes.

The SCP approach in this study has proven to be instrumental in revealing dynamic, intercellular differences and linking individual cellular proteomes with the status of symbiosis. Compared with single-cell RNA sequencing (sc-RNAseq) methods, which can also provide a quantitative view of the gene expression and interactions of microbial eukaryotes ^47–49^, proteomic pipelines cannot amplify molecular signals and are thus more dependent on protein abundance in individual cells and on the sensitivity of DIA-MS platforms. However, SCP has the following advantages over sc-RNAseq. Proteins are generally the molecules that perform functions or form cellular components, in contrast to mRNA molecules that still need to be translated. Moreover, many studies have shown that the correlation between protein and mRNA abundance is only modest ^50,51^, suggesting that protein abundance is a more direct indicator of cellular physiology. While proteins or their peptide fragments cannot be amplified for signal detection, the absence of this step also prevents any potential amplification biases toward specific molecules, which can be introduced during library preparation for scRNA-seq. For example, nucleic acids with high-and low-GC content tend to have lower sequencing yields ^52^, which may be the case for *P. bursaria* with its GC content of only 28.8% ^53^. Proteins, and their digested peptide fragments, are also much smaller than their corresponding mRNA molecules, which can be more difficult to extract for cells such as *Chlorella* with rigid cell walls that are not easy to fully break down ^54^. Finally, we coupled proteomic analyses with a large-particle sorter that uses air streams to divert droplets, which can be gentler than hand picking that can cause mechanical shock to *P. bursaria* and trigger trichocyst discharge and other proteomic changes. Our method further minimizes sample loss by avoiding liquid transfer and using low-binding Evotips for direct sample loading for liquid chromatography. As an alternative to scRNA-seq, this SCP protocol can be applied to investigating other interactions among microbial eukaryotes.

## Materials and Methods

### Cell culturing

*P. bursaria* DK2 cells were maintained with 1 mg/mL penicillin and 200 ug/mL kanamycin and fed with axenic cultures of the non-symbiotic food alga *Chlorogonium capillatum* (Chlamydomonadales, Chlorophyceae; strain NIES-3374, National Institute for Environmental Studies, Japan) every 3-4 days. *P. bursaria* DK2 and *C. capillatum* were cultured in sterile Modified Chlorogonium medium (CGM) of 0.015% (w/v) sodium acetate, 0.075% (w/v) yeast extract, Dryl’s buffer (2 mM sodium citrate, 1 mM NaH_2_PO_4_/Na_2_HPO_4_ and 1.5 mM CaCl_2_) and Volvic mineral water with a 12:12 light-dark cycle at 23℃. Chlorella-symbiotic *P. bursaria* cells were cultivated under a consistent light condition (photosynthetic photon flux density (PPFD): 21.8 μmol·m^−2^·s^−1^). Aposymbiotic cells were generated from the symbiotic cells by treating them with 100 μg/ml of paraquat dichloride ^55^ (Cat #856177, Sigma-Aldrich, MA, USA) for approximately one week in a light-to-dark cycle of 12 h:12 h and maintained under a lower light condition (PPFD: 6 μmol·m^−2^·s^−1^).

### Endosymbiosis establishment

Aposymbiotic DK2 cells and *Chlorella* cells were prepared for the reinfection experiment. Five days before the re-infection, aposymbiotic DK2 cells were fed with *C. capillatum*. Two days later, the remaining *C. capillatum* were removed along with any cell debris by filtering with a 10-µm nylon mesh and washing with Dryl’s buffer. *Chlorella* cells were extracted from stable symbiotic DK2 cells with the following procedure. First, symbiotic DK2 culture was filtered through eight to twelve layers of gauze. Then the cell debris was filtered away with 10-µm nylon mesh, and the remaining symbiotic DK2 cells were washed with Dryl’s buffer for three times. These cells were resuspended in Dryl’s buffer and centrifuged by Heraeus Multifuge X3R centrifuge (Thermo scientific, U.S.A) with 20,000 RCF for 10 minutes to break the cells. The released *Chlorella* cells were resuspended in 1X PBS buffer and centrifuged by Heraeus Fresco 21 centrifuge (Thermo scientific, U.S.A) at 5,000 rpm for 1 minute for three times to wash away bacteria and cell debris. After resuspending *Chlorella* in Dryl’s buffer, the cell density was measured by Cytoflex S (Beckman Coulter, United States). The density of the starved aposymbiotic DK2 cells was determined under the microscope ZEISS Axio Lab.A1 (Zeiss, Jena, Germany). For the re-infection, aposymbiotic *P. bursaria* was mixed with *Chlorella* cells at a ratio of 1:10000 to a final culture volume of 53 mL and incubated at 23℃ with a 12:12 light-dark cycle and a PPFD of 6 μmol·m^−2^·s^−1^. After 1 hour of incubation, the non-engulfed free-floating *Chlorella* cells in the culture were removed by filtering with a 10-µm nylon mesh and washing with 1X Dryl’s buffer three times. The *P. bursaria* cells were resuspended in CGM and put back into the growth chamber. At 1 DPI, light intensity was raised to a PPFD of 21.8 μmolm^−2^s^−1^, which is the same as the condition for culturing symbiotic *P. bursaria* cells. The non-symbiotic food alga *C. capillatum* was fed to *P. bursaria* at 1, 4, 7, 14, and 21 DPI after sampling cells for flow cytometry and microscopy.

### Time-series sampling and large-particle single-cell sorting

At each time point, 1.5 mL of culture was sampled, including 0.5 mL fixed with 4% formaldehyde for microscopy analyses and 1 mL for flow cytometry and sorting. We used a large-particle flow cytometer and sorter COPAS Vision (Union Biometrica Inc., Massachusetts, USA) equipped with a 1000-µm flow cell and a 488-nm laser. For fluorescence-activated cell sorting, DK2 cells were diluted 40X with ddH_2_O, transferred to COPAS Vision, selected by gating particles with time of flight at 560-1040 (corresponding to ∼75-140 µm) and extinction at 320-760 a.u., and were sorted into V-bottom tubes (Scientific Specialties, Inc., Lodi, CA, U.S.A) already coated with a non-ionic detergent n-dodecyl-β-D-maltoside (DDM). In addition to the brightfield image and the cell size (time of flight), chlorophyll a fluorescence of each sorted cell was measured by COPAS VISION 1000 (Union Biometrica, USA) during cell isolation based on the autofluorescence of algae inside DK2 cells using a far-red fluorescence detector (680/42 nm) with 488-nm laser excitation. All flow cytometry and sorting analyses had consistent gains and PMT voltage settings. Sorted DK2 cells were randomly selected for subsequent LC-MS/MS analysis.

### Microscopy analyses

For each time point, the 0.5-mL fixed sample was examined using the phase contrast module of ZEISS Axio Lab.A1 (Zeiss, Jena, Germany), with fluorescence imaging using a mercury lamp with a 590-nm emission filter to visualize autofluorescence of algae. The fluorescence levels were measured using ImageJ. For samples with high endosymbiotic levels (14 and 21 DPI), cells were scanned under a confocal microscope ZEISS LSM880 (Zeiss, Jena, Germany), with excitation at 488 nm and a GaAsP detector (emission range: 620-729 nm). Z-stack images were integrated into 3-D graphics and cyclized fluorescence of algae was calculated with IMARIS 9.5.1(Oxford Instruments, United Kingdom).

### Single-cell proteomics sample preparation

Cells were lysed following a previous protocol ^56^ by adding 2 µL of lysis buffer (1% DDM / 50 mM triethylammonium bicarbonate (TEAB)) to each V-bottom tube containing one sorted *P. bursaria* cell. The tubes were sonicated in a water bath (Ultrasonic Cleaner CPX3800H/Branson) for 10 cycles of 30 seconds on and 30 seconds off. Following sonication, the tubes were centrifuged at 3000g for 5 minutes at 4 °C. Protein digestion was initiated by adding 1 µL of Lys-C stock (5×10⁻⁵ I.U./µL) to the sample. Samples were incubated at 37 °C for 1 hour. Subsequently, 1 µL of trypsin stock (50 ng/µL) was added, and the samples were incubated overnight at 37 °C. After the overnight digestion, the tubes were centrifuged at 3000 g for 5 minutes at 4 °C. The reaction was quenched and acidified by adding 2 µL of 5% formic acid. The samples were centrifuged again at 3000g for 5 minutes at 4 °C. The supernatant from each sample was transferred to an Evotip (Evosep Biosystems, Odense, Denmark), which minimizes loss during subsequent sample loading.

### Single-cell LC-MS/MS

LC–MS/MS analysis was performed using an Evosep One liquid chromatography system (Evosep Biosystems, Odense, Denmark) coupled to a timsTOF HT mass spectrometer (Bruker Daltonics, Bremen, Germany). To avoid potential batch effects, all the single-cell samples from all the time points were analyzed in a single batch. Samples were loaded onto Evotips, and peptides were separated using the Whisper 40 samples-per-day method on an EV1112 Performance column maintained at 40 °C and interfaced with a CaptiveSpray ion source. Data were acquired in DIA-PASEF mode over an m/z range of 400–1,200 Da and an ion mobility range of 0.6–1.6 1/K₀, using 16 × 25 Da isolation windows and two TIMS ramps per cycle. Accumulation and ramp times were both set to 100 ms, yielding a total cycle time of 1.8 s.

### Proteomics data processing

Raw data were processed in Spectronaut v. 19.0.240606.62635 (Biognosys AG, Switzerland) using the “directDIA” workflow with default settings. Database searches were performed against a merged database, Merge_PbCvCgPBCV1, combined protein sequences predicted from *P. bursaria* Dk2 whole-genome sequences (GCA_043974915.1; GCA_043974975.1), *C. variabilis* NC64A nuclear genome (JGI project ID: 16663), *C. variabilis* NC64A mitochondrial genome (NC_025413.1), *C. variabilis* NC64A chloroplast genome (NC_015359.1), *Paramecium bursaria* Chlorella Virus 1 (PBCV-1; NC_000852.5), and *C. capillatum* (Supplementary Table S3). Trypsin was specified as the digestion enzyme, allowing up to two missed cleavages. Carbamidomethylation (Cys) was set as a fixed modification, while oxidation (Met) and acetylation (protein N-terminus, Lys) were set as variable modifications. False discovery rates (FDRs) for peptide-spectrum matches, peptides, and proteins were controlled at 1% using scrambled decoy sequences, with the dynamic decoy size set to 10% of the library size. Quantification was based on MS2 fragment ion intensities with local cross-run normalization enabled. Peptide precursor abundances were determined by summing the peak areas of fragment ions, and peptides were grouped by modified sequences. Protein-level quantification was performed using standard spectral library settings, and peptide signal intensities were summarized as the mean precursor area for reporting.

### Data analyses

Out of the 80 *P. bursaria* SCP samples, we filtered out 4 that had < 150 protein groups. Protein groups that are present in < 3 cells were also excluded from the downstream analyses. SCP data were analyzed in R v. 4.5.1 and using the package Seurat v. 5.4.0 ^57^. Flow cytometry and microscopy-derived data were visualized using ggplot2 v. 4.0.1. The differences in protein abundance among samples were assessed by one-way ANOVA using ‘stats’ v. 4.5.1, followed by Tukey’s HSD and pairwise comparisons using ‘emmeans’ v. 2.0.1 in R. For heatmap visualization in Figs. 3-5, protein abundances in each cell were normalized using Seurat’s LogNormalize method, scaled to 1,000,000 per cell, and transformed by the natural logarithm (log1p). Heatmaps were generated from z-score–normalized data (per protein group) with ‘pheatmap’ v. 1.1.13 in R. Highly variable protein groups were selected using the mean variance plot (MVP) method and scaled, followed by PCA dimensionality reduction. For UMAP visualization of cellular proteomes, nearest neighbor graph and clustering were performed using the top 5 PCs. Candidate proteins with lower abundance at 2 DPI were isolated by Wilcox rank-sum test (adj.p < 0.05, avg log_2_FC < -1). Differentially abundant proteins between high- and low-chl cells at 14 DPI or 7 and 14 DPI were detected using Wilcox rank-sum test (p < 0.1, avg log_2_FC > 2). R coding was assisted by AI tools: Claude v. 4 and Perplexity (Grok 4.1). For phylogenetic analyses of SLC proteins, we aligned the amino acid sequences using MAFFT v7.526 ^58^ and reconstructed maximum likelihood trees using IQ-TREE v2 ^59^ under the best-fit model Q.yeast+F+R7 selected by ModelFinder ^60^. Phylogenies were visualized using Interactive Tree of Life (iTOL) v6 ^61^.

## Data availability

The MS proteomics data have been deposited to the ProteomeXchange Consortium via the PRIDE ^62^ partner repository with the identifier PXD073244.

## Acknowledgements

We thank the technical support provided by the Comprehensive Flow Cytometry Laboratory and the IPMB Cell Biology Core Lab Division Light Microscopy, Academia Sinica, Taiwan. We are grateful to Ming-Wei Lai, Jeremy Catinot, and Yu-chi Liu for laboratory support and discussion. This work was funded by Academia Sinica grants AS-CDA-110-L01 (C.K.), AS-IA-110-L01, and AS-GCS-113-L03 (J.-Y.L.) and National Science and Technology Council (Taiwan) grants 111-2611-M-001-008-MY3, 114-2628-B-001-012- (C.K.), 113-2326-B-001-002 (J.-Y.L.), and 112-2311-B-001-024-MY3 (C.-C.H). M.M.K. was supported by an NSTC postdoctoral fellowship (NSTC 113-2811-B-001-065).

## Supplementary Tables

Table S1. List of isolated single *Paramecium* cells

Table S2. Abundance of detected protein groups in each *Paramecium* cell

Table S3. Sequence databases for single-cell proteomic analyses

Table S4. Abundance of detected trichocyst matrix proteins (TMPs) in each *Paramecium* cell

Table S5. List of 2DPI-lower proteins of *P*. *bursaria*

Table S6. The differentially abundant proteins between high- and low-chl *P*. *bursaria* cells at 7 and 14 DPI

Table S7. Homologs, allelic variation, and predicted localization of the SLC protein in *P*. *bursaria* DK2

Table S8. List of *Chlorella* proteins and their functional annotations

## Extended Data Figures

**Extended Data Figure 1.**
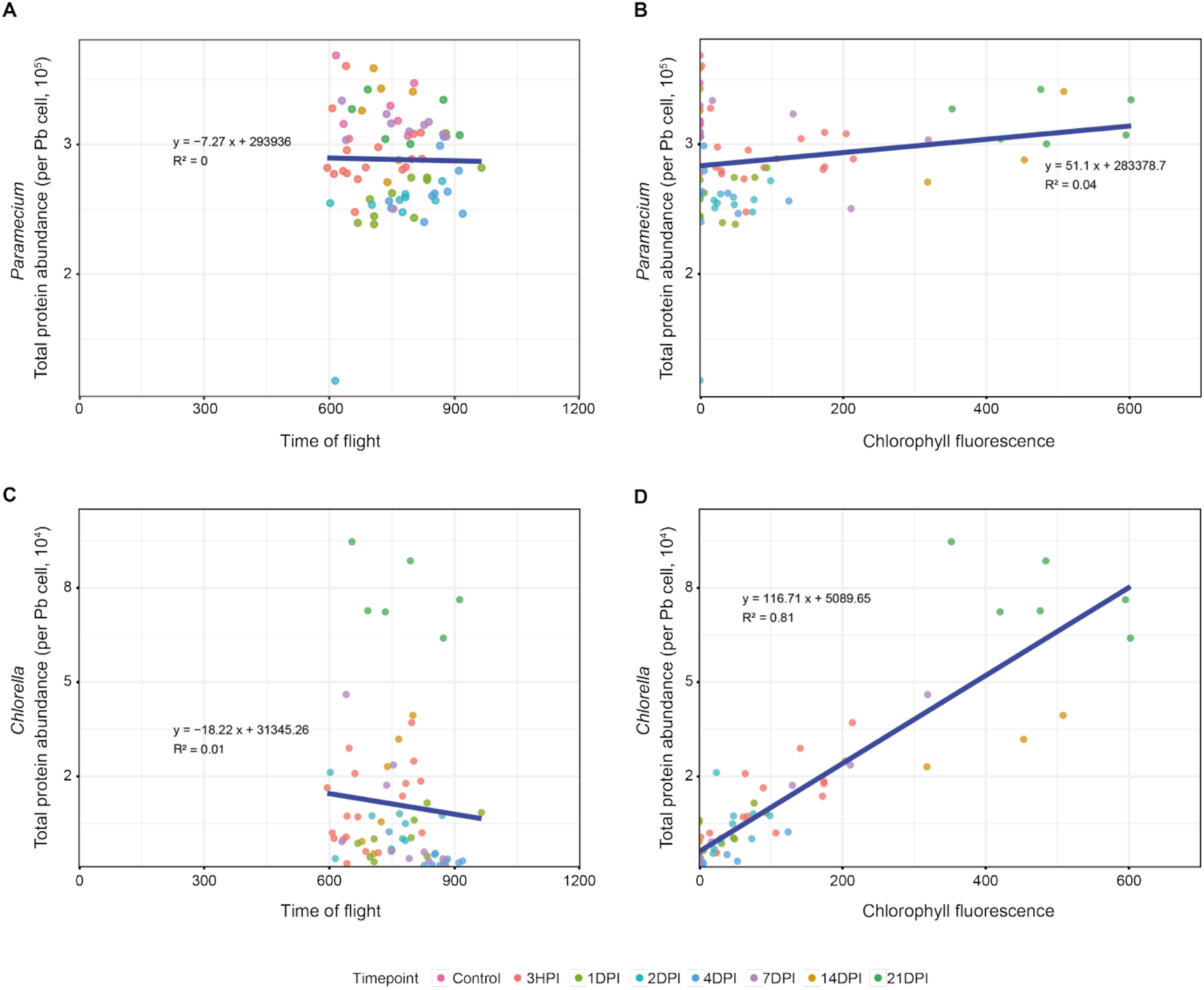
Total protein abundance, time of flight, and chlorophyll fluorescence of individual *P*. *bursaria* cells. The total abundance of detected *Paramecium* and *Chlorella* proteins is plotted against the time of flight (560-1040, roughly corresponding to a cell size of 75-140 µm; A and C) or chlorophyll fluorescence intensity (B and D) of single *P*. *bursaria* cells at different timepoints. A positive correlation (R^2^ = 0.81) can be observed between the total *Chlorella* protein abundance and chlorophyll fluorescence.

**Extended Data Figure 2.**
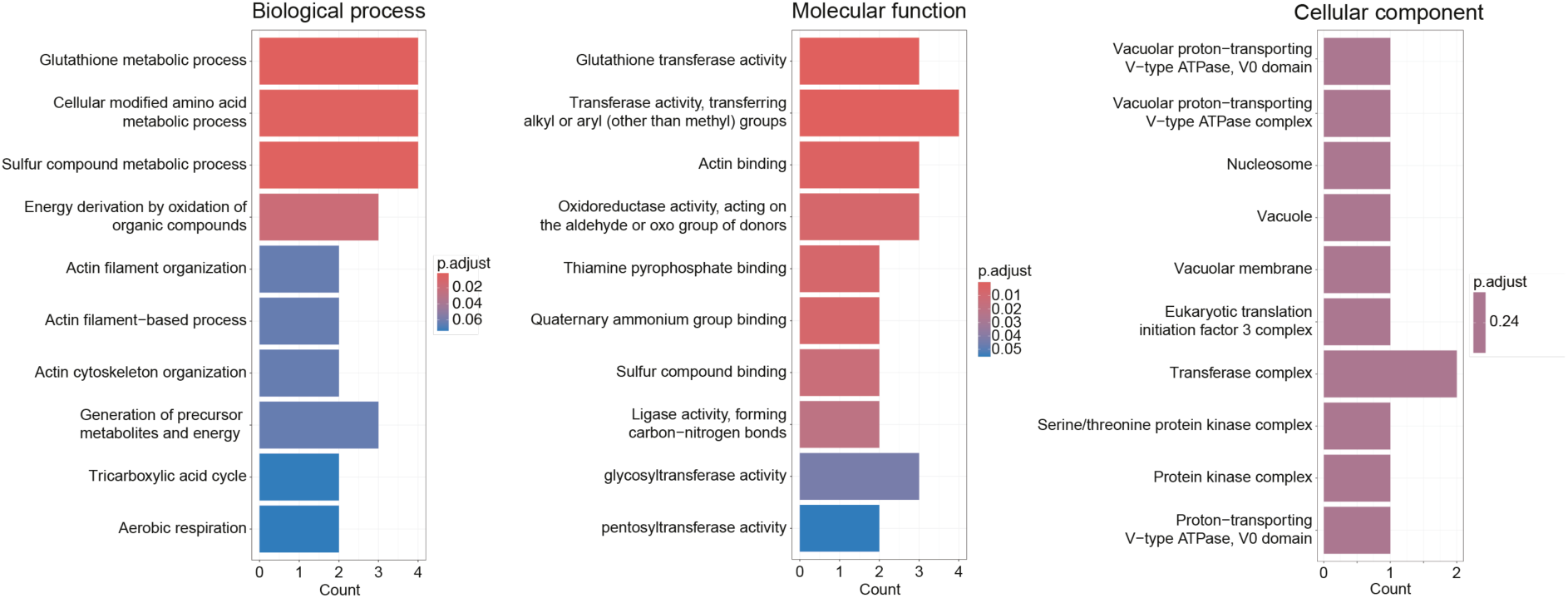
GO functional enrichment analysis of the *P*. *bursaria* proteins with lower levels at 2 DPI. The top enriched Gene Ontology terms of the 2DPI-lower proteins (compared against all detected *P*. *bursaria* proteins) are shown for each of three major categories.

**Extended Data Figure 3.**
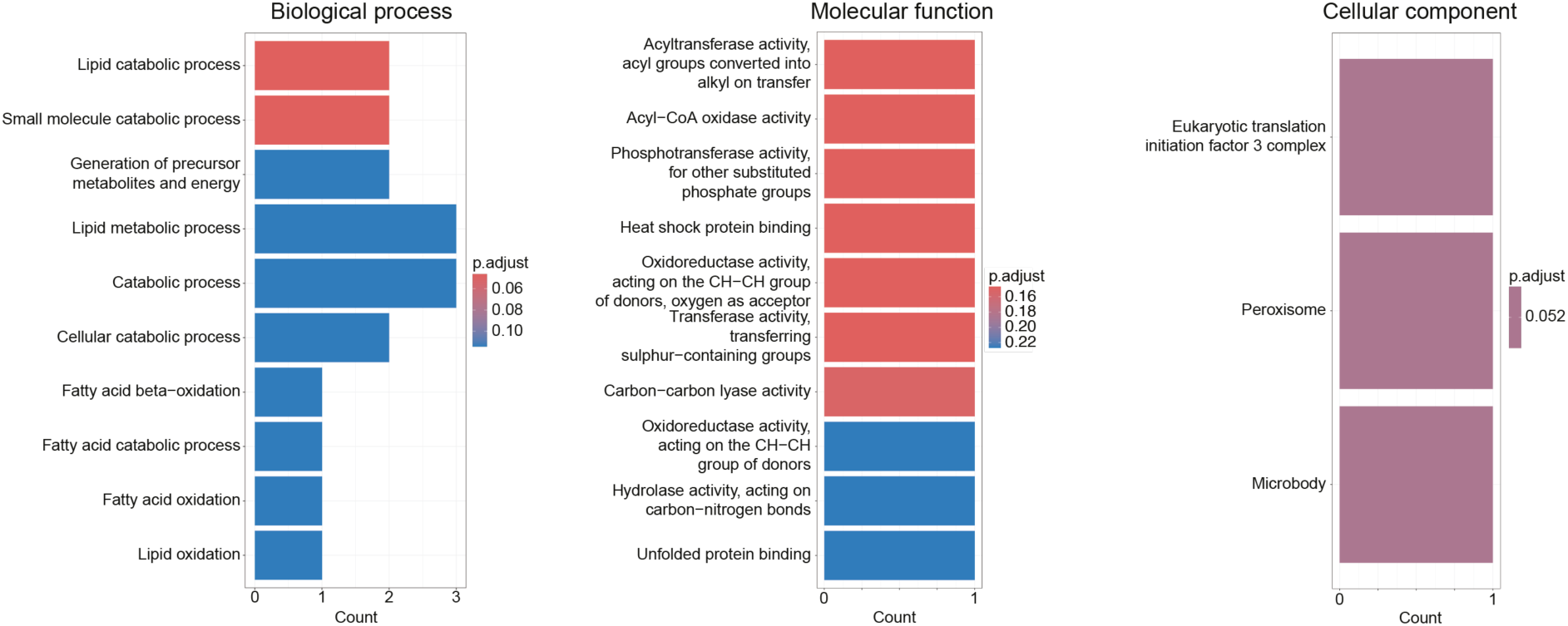
GO functional enrichment analysis of the differentially abundant proteins between high-chl and low-chl *P*. *bursaria* cells at 7 and 14 DPI. The top enriched Gene Ontology terms of the *P*. *bursaria* proteins that are differentially abundant between high- and low-chl *P*. *bursaria* cells at either 14 DPI or both 7 and 14 DPI (Fig. 4C). The enrichment analysis was conducted by comparing these proteins against all detected *P*. *bursaria* proteins.

**Extended Data Figure 4.**
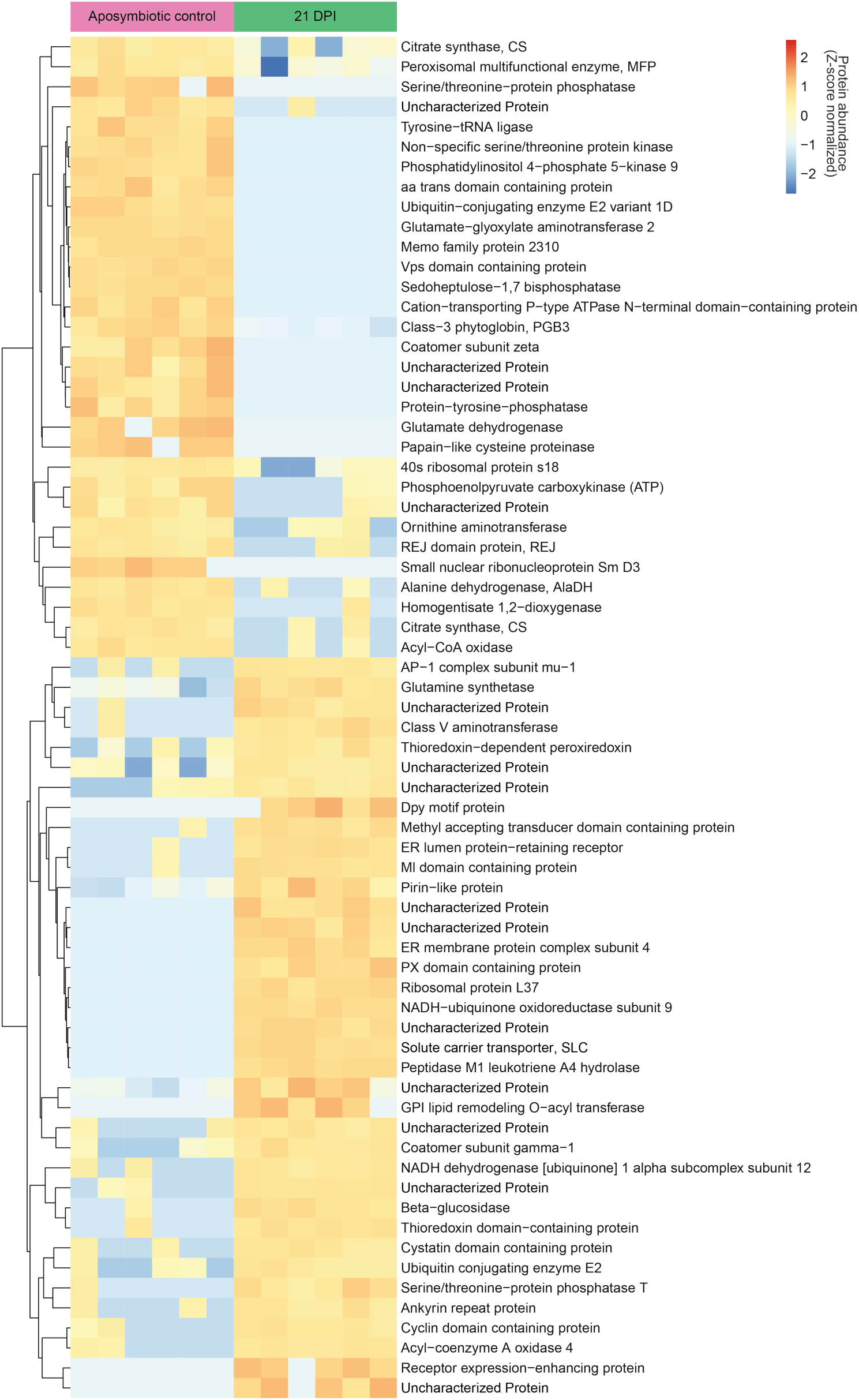
Comparison between 21-DPI (stable symbiotic) and aposymbiotic *P*. *bursaria* cells. Normalized abundance profiles and annotations are shown for 68 *P*. *bursaria* proteins differentially abundant (fold change > 4 or < -4; Wilcoxon rank-sum test: *p* < 0.01) between 21-DPI (all high-chl) and aposymbiotic control cells.

**Extended Data Figure 5.**
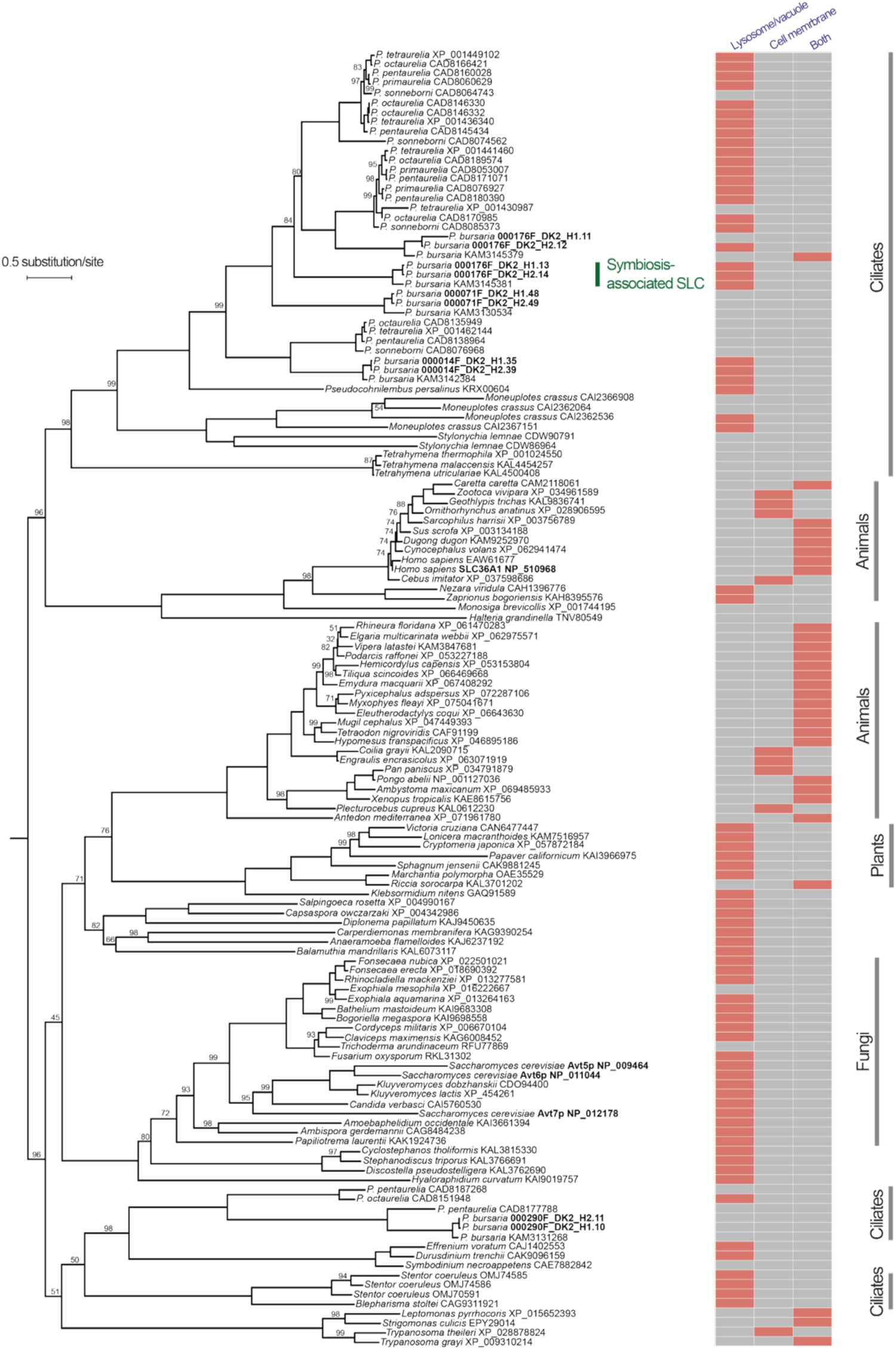
Phylogeny and predicted localization of the symbiosis-associated SLC and related proteins in eukaryotes. The maximum likelihood tree is mid-point rooted, with percentages of bootstrap support (if lower than 100) labeled for internal branches. DeepLoc 2.0 predicted localization (if the probability is over the threshold; Table S7) is shown for each sequence.

## Notes

### Competing Interest Statement

The authors have declared no competing interest.

